# An Unsupervised Learning Method for Disease Classification Based on DNA Methylation Signatures

**DOI:** 10.1101/492926

**Authors:** Mohammad Firouzi, Andrei Turinsky, Sanaa Choufani, Michelle T. Siu, Rosanna Weksberg, Michael Brudno

## Abstract

Recent work has shown that genome-wide DNA methylation (DNAm) profiles can be used to discern signatures that can identify specific genetic disorders. These methods are especially effective at identifying single gene (Mendelian) disease, and methods to identify such signatures have been built by comparing methylation profiles of known disease versus control samples. These methods, however, have to-date been supervised, precluding the application of these methods to diseases with as-yet-unknown genetic cause. In this work, we tackle the problem of unsupervised disease classification based on DNAm signatures. Our method combines pre-filtration of the data to identify most promising methylation sites, clustering to identify co-varying sites, and an iterative method to further refine the signatures to build an effective clustering framework. We validate the proposed method on four diseases with known DNAm signatures (CHARGE, Kabuki, Sotos, and Weaver syndromes) and show high accuracy at determining the correct disease using unsupervised analysis. We also experiment with our approach on a novel dataset of patients with a clinical diagnosis of Autism, and illustrate the de novo identification of a specific subtype.

## 1 Introduction

Epigenetic studies have shown existence of a close relation between epigenetic marks over the DNA, such as DNA methylation (DNAm), and human disorders [1]. Among epigenetic marks, DNA methylation, characterized by the presence of a methyl group at a CpG dinucleotide is one of the most broadly interrogated, both because of technology that can provide genome-wide profiling of methylation at CpG sites, such as methylation arrays (Illumina 450k and Epic) and sequencing-based protocols, such as reduced representation bisulfite sequencing (RRBS), and the close relationship between methylation in promoters of genes and gene expression. Recently DNAm signatures have been suggested as a mechanism to differentiate between clinically similar rare disorders and identify their genetic causes. Because the pathogenesis of these disorders involves a single gene or protein, each disease has a specific effect on methylation of various regions of the human genome. Several recent studies have addressed the use of DNAm data to extract disease signatures as a supervised learning problem [2, 3, 4, 5]. It is hoped that the signatures can be used in conjunction with more standard clinical testing (e.g. exomes) to resolve variants with unclear clinical significance (VUS’s) or identify when a disease causative variant was potentially missed (e.g. due to being outside of coding regions of the gene).

Generally, the approaches that have been used in previous studies to elucidate the DNAm signature for a disease is to run statistical tests on training data consisting of samples with particular disorders and a number of healthy samples (controls) to extract the differentially methylated regions (DMRs). Each DMR typically contains one CpG site with statistically different methylation profile between cases and controls, as characterized by a statistical test. These DMRs are then used to train a supervised model, such as an SVM, to classify disease and healthy samples.

This problem was studied in [3] for differentiating Sotos and Weaver syndromes. These two disorders are clinically similar, however they are caused by mutations in two separate genes (*NSD1* versus *EZH2*, respectively). The model was trained using known harmful (truncating) mutations, and validated both on a held-out set and on mutations without a deifnitive explanation (missense mutations labelled as “Variants of Unknown Significance”, or VUS, on a clinical report). After identification of specific CpG sites as DMRs, a hierarchical clustering model was used to identify the signature.

A similar approach was then applied in [2] for two more syndromes, Charge and Kabuki, and a group of controls. *β*-values [6] were extracted from methylation values (M-values) and supervised statistical tests were used to detect DMRs. Using DMRs, a support vector machine (SVM) model was trained with a linear kernel for each disease cohort and a matching set of controls. The predictive models for each syndrome were also tested on the other syndrome samples to show the specificity power of the trained models. A similar supervised learning approach to [2] was used in [4] where a multi-class SVM model was trained to distinguish between multiple syndromes using genome-wide *β*-values. To detect DMRs, a bump hunting approach was used [7].

In this work we tackle the problem of identifying disease samples using DNAm array data in an unsupervised framework, which to our knowledge has not been studied previously. The unsupervised setting for this problem is important because there are more than 7, 000 rare disorders, while DNAm signatures for only a handful are known. Unsupervised approaches are especially useful for looking at more common disorders, such as Autism, which are known to include many single-gene disorders with overlapping phenotypes [8]. The problem of (unsupervised) clustering of DNAm profiles is particularly challenging not only because of its high-dimensional nature (we utilize 450k methylation arrays, so have 450,000 features for every sample) and batch effect issues, but also because unlike supervised approaches, we cannot extract DMRs with simple statistical tests.

We devise an unsupervised procedure and apply it to DNA methylation array data consisting of patients with specific neuro-developmental disorders studied by [2] and [3]. We also apply our method to a new data set with a collection of autism spectrum disorder (ASD) samples and controls. Authors in [9] conducted a histone-acetylome-wide association study (HAWAS) and extracted differentially acetylated locations for different brain regions of samples affected by specific variants of ASD in a supervised framework. Using our unsupervised method, we identify a sub-cluster of ASD samples using our method with consistent DNAm signatures. Overall our method shows high accuracy at delineating multiple neurodevelopmental disorders without any *a priori* of patients with a specific disease.

## 2 Background

In this section we provide the key machine learning concepts that are relevant to the rest of the paper.

Denote the data with 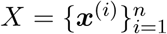 of *n* independent samples, where each ***x***^(*i*)^ is a *d* dimensional vector in ℝ^*d*^. The aim is to find a partitioning of *X* into a fixed number of non-empty disjoint groups or clusters *C*_1_, *C*_2_, *…, C*_*K*_ where *k* represents the number of clusters. We also indicate the set of data features or variables by *F* = *{f*_1_, *f*_2_, *…, f*_*d*_}.

### 2.1 Clustering

Searching for the clusters *C*_1_, *C*_2_, *…, C*_*K*_ based on specific criteria is a broad area of study and is not the focus of this work [10]. We briefly discuss simple methods that are widely used in genomic studies and also can be aligned with our proposed approach.

k-means clustering algorithm uses the Euclidean distance as the dissimilarity measure between the data points. *C*_1_, *C*_2_, *…, C*_*K*_ are founded so that 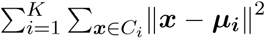 is minimized. Each ***µ***_***i***_ indicates the mean of the members of *C*_*i*_.

In bottom-up hierarchical clustering methods, each sample starts with its own cluster. At each step, pairs of most similar clusters are merged to ultimately achieve to final clusters *C*_1_, *…, C*_*K*_ using the chosen similarity measure.

Gaussian mixture model (GMM) is a type of latent probabilistic models in which the distribution of samples given the *j*-th cluster is modeled as a Gaussian distribution *𝒩*(*µ*_*j*_, ∑_*j*_) where *µ*_*j*_ and ∑_*j*_ are the Gaussian’s mean and covariance matrices. Denoting all the model parameters with *θ*, the marginal likelihood function *ℒ*(*X |θ*) is then maximized using the expectation-maximization (EM) algorithm. The values of the latent components specifies the cluster memberships.

#### 2.1.1 Bi-clustering

Bi-clustering methods (also known as co-clustering) have been applied to DNA gene expression data to extract subgroups of genes that are co-expressed across only a subset of samples and may indicate biological implications. The closest bi-clustering category to our method is the two-way clustering (TWC). These methods are based on alternatively performing clustering over genes, and samples until particular criteria are met. Bi-clustering methods work well in practice in cases where the contamination of the samples is limited, and the samples are well defined and well replicated [11]. This fact limits the use of bi-clustering methods for DNAm array datasets, which generally include only a few samples and are impacted by various type of noise.

A difference between our proposed method and TWC bi-clustering approaches is that the latter directly uses the features and samples to detect co-expression, but our proposed framework extracts low dimensional signals from a group of features and uses them to cluster the samples.

### 2.2 Feature Selection

To avoid the curse of dimensionality, high computation complexities, and possible overfitting for high-dimensional data with a small sample size, a natural step is feature selection. Various feature selection methods have been proposed and are generally classified into three types: filter-based, wrapper, or embedded methods [12]. In literature, feature selection is often based on filtering methods in which a subset 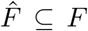 is chosen; generally, 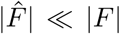. For example, in supervised learning settings, features are often selected by their mutual information with the classes [13], or based on P-values from statistical tests such as Student’s t-test and Mann-Whitney-Wilcoxon-test [14].

Feature selection becomes more challenging when class labels are not available. A simple unsupervised approach is variance filtering, which has been broadly used before [15]. A diverse set of unsupervised filtering methods have been proposed [16]. While our focus in this work is not to compare these methods, Appendix B presents the potential negative effect of arbitrarily choosing the feature selection approach on DNAm array data. Briefly, since batch effect is a critical problem for DNAm array data, commonly used approaches such as nearest neighbor graph-based filtering methods might cause the selected features to be biased toward discriminating batches rather than the underlying signals. In [17], authors propose a batch effect removal approach for single-cell RNA sequencing data using mutual nearest neighbor data points. Based on our experiments, we suspect that a similar approach might be applicable for DNAm array data as well.

### 2.3 Representation Learning

Representation learning is nowadays tied with deep architectures to learn multiple levels of representation, or a hierarchy of features [18]. The challenges of using deep architectures for DNAm array data is further explained in the discussion section. In this work, we briefly explain two traditional approaches, principal component analysis (PCA) and independent component analysis (ICA) which are simple and popular dimension reduction methods for microarray data.

Linear PCA gives a representation of data that projects the samples on a set of orthogonal principal components. Principal components are the singular eigenvectors of the zero centered data matrix. To obtain a *d*^*′*^ *≤ d* dimensional representation, the first *d*^*′*^ principal components are chosen. Intuitively, projection of the data on these vectors preserves the maximum fraction of data variance.

Linear ICA assumes that a data sample ***x*** is generated from a vector of hidden signals ***s*** = (*s*_1_, *…, s*_*d′*_) by a linear transformation *A* as ***x*** = *A****s***. The task is then to learn *A* and ***s*** in a way that components of ***s*** are maximally independent according to a measure of independence. Two popular notions of independence are mutual information, and non-Gaussianity.

## 3 Proposed Method

In this section, a more general framework of the our proposed method, FCRC, is described. The name of the proposed method is based on its steps (Filtering features, Clustering variables, Reducing the clusters and the dimension, and Clustering samples). An overview of FCRC is shown in Algorithm 1, and the details as implemented for our experiments is provided in Appendix A. FCRC stages are visualized in Figure 1. Further details and the hyperparameter values are provided in the experimental results.

**Figure 1:**
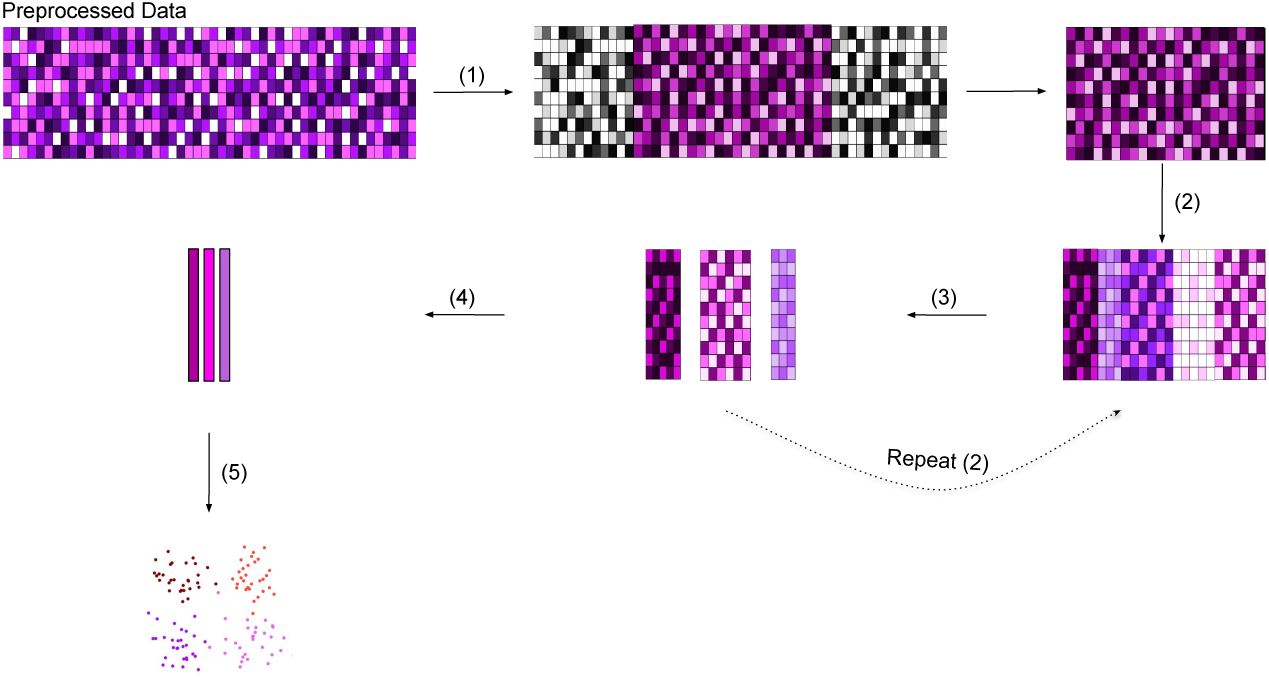
FCRC stages for DNAm array data: (1) A subset of variables with the highest IQR values are selected. (2) Selected variables are clustered based on a similarity measure for variables (3) Redundant variable clusters are dropped based on specific statistical criteria (4) A dimensional reduction approach is used to obtain a low dimensional representation vectors based on variables within each variable cluster (5) Samples are clustered using the final representations.

### 3.1 Preprocessing

A careful selection of preprocessing steps can reduce variance and thus improve statistical power of processing and analyzing DNAm data [19]. In particular, several useful and popular preprocessing methods (e.g. data normalization) are implemented in minfi, a Bioconductor package in R [20]. The initial step in our framework is a potential preprocessing that may improve the downstream analysis performance of the data. To reduce the batch effect problem, since the data generally contains relatively small sample size in each batch, a popular method for adjusting known batches based on empirical Bayes frameworks is used [21].

### 3.2 Filtering Features

We apply an unsupervised filtering-based feature selection using the inter-quantile ranges (IQRs) of each *f*_*i*_ and selected 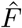 as the union of features with the highest IQR values. In our experiments, this approach effectively removes the redundant features.

### 3.3 Clustering Variables and Dimension Reduction

In the FCRC, feature selection in the previous step may be insufficient to lower the data dimensionality significantly. To further reduce the potential overfitting problem and computational cost, for a predetermined number *L ≥* 2, we partition 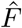 into *L* clusters as 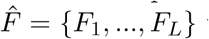 where features in each subset *F*_*i*_ have high similarity, with the hope that each cluster variables are closely related and have a similar information. Feature clustering is implemented in ClustOfVar package in R [22] using Pearson correlation as the measure of similarity among features. For each *l ∈* {1, …, *L*}, we then extract a low dimensional representation vector 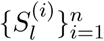 for the samples based on only the features in cluster *F*_*l*_. In our experiments, PCA and ICA are used.

Accurate unsupervised detection of disease groups may be impeded by several factors, including outliers within each disease group and batch effects. To handle both problems, for each *l ∈* {1, …, *L*}, we clustered individuals into two groups using the variables in *F*_*l*_. For *F*_*l*_ to be useful we impose the constraint that it should not result in a clustering with less than 4 samples in a group. In such cases we drop the corresponding cluster.

#### Algorithm 1 FCRC procedure

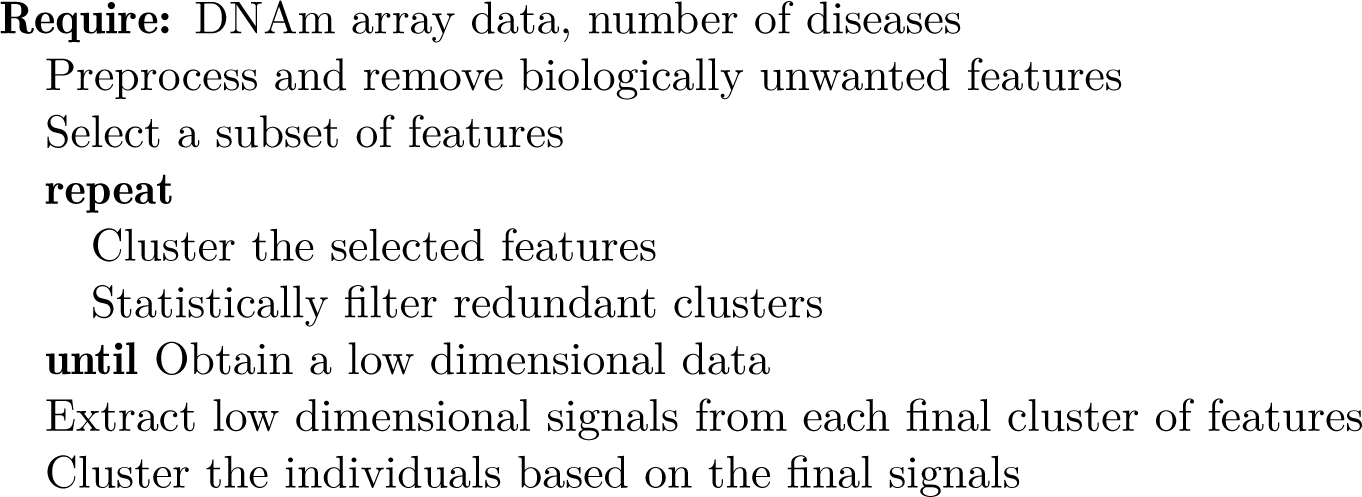

While we perform batch correction before clustering, to tackle any residual batch effects we define an *absorption ratio* for each variable cluster. We look at the Kolmogrov-Smirnov nonparametric statistical test score for the shift between values of that feature over the two groups of individuals that were obtained from each *F*_*l*_, and assign that feature to the variable cluster which has the highest test score. A feature might not get absorbed by any clusters if the shift between the mean values of that feature over the two groups of each variable cluster is not significant enough. The absorption ratio of *F*_*l*_ is then defined as the proportion of features that are absorbed by *F*_*l*_. Our assumption here is that a batch effect causes a shift in a large proportion of data features genome-wide (i.e. high absorption ratio), whereas a disease affects a small portion of DNA methylation values over genome (lower absorption ratio). Therefore, we eliminate the variable clusters that have an absorption score above a constant. We denote the set of remaining variable clusters with 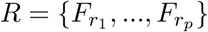 where *p ≠ L*. To get the final data, we concatenate the corresponding signal vectors as 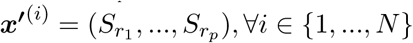.

We note that the process of feature clustering and removing the redundant signals might be done multiple times until the data dimensionality becomes sufficiently low. In our experiments, this process is done just once.

### 3.4 Final Sample Clustering

The final stage is to divide the data points into *K* groups presumed to represent different disorders. We cluster the samples using 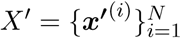 to get the final clusters. In our experiments, we used simple clustering methods like Gaussian mixture model (GMM) with the expectation-maximization (EM) algorithm for optimization and k-means clustering.

## 4 Experimental Results

We demonstrate the performance of our method on several DNAm array datasets. To validate the ability of our method to reconstruct DNAm signatures for known diseases we apply FCRC to data from [3] and [2], which contains 39 samples with CHARGE, 19 samples with Kabuki, 38 samples with Sotos and 7 samples with Weaver syndromes, as well as 90 controls. We initially applied the method separately on CHARGE/Kabuki and Sotos/Weaver, reflecting the previous work, and clustered each of these into 3 clusters (two disorder clusters and the controls). We subsequently combined these datasets to create a more challenging clustering problem. In this case the samples are clustered into into five groups: one group for each disease, and one for all the controls in the two data. Well known clustering quality measures are computed to evaluate the performance of the FCRC on these clustering tasks.

Given the strong performance of FCRC on the initial data, we applied it to a novel dataset containing different types of autism syndrome patients (ASD), with the task of distinguishing particular subtypes from a group of known ASDs and their controls.

### 4.1 Clustering Quality Measures

In our work we utilize three different clustering performance measures to quantify the results. Given a set of clusters *S*, set of classes *C*, both partitioning *n* data points, the Purity is computed as

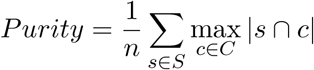

Purity does not perform well for imbalanced data. To ensure that this will not affect the clustering performance two other clustering measures are also calculated.

Another measure that is used is the normalized mutual information (NMI) which is defined as

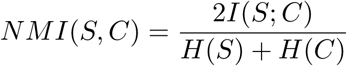

where *I* and *H* represent the mutual information and the entropy, respectively. The value of NMI is robust to the increase of cluster numbers.

Rand index is another measure which calculates the proportion of pairs that are correctly clustered in the same or different groups. Adjusted Rand index (ARI) is corrected-for-chance version of the Rand index. The ARI value can vary between −1 and +1. Higher Purity, NMI, or ARI values indicate better clustering performance. The confusion matrices of the clustering results are also reported in Appendix D.

### 4.2 CHARGE and Kabuki Syndromes Data

In this section we report the results on the CHARGE and Kabuki data [2]. The data consists of two groups: discovery cohort, and validation cohort, including patients with CHARGE and Kabuki syndromes, and a number of controls. We did our experiments on both the discovery cohort and the union of both discovery and validation cohorts. In addition to pathogenic samples and controls, as described in [2], there are variants without uncertain significance (VUS). These samples are excluded from our clustering performance analysis. More details about the samples and their population is provided in the [2].

In this (and all other experiments) we utilize the M-values extracted from the samples’ raw meta-array data. IQR filtering is used to choose 2.0 – 5.0 percent of variables with the highest IQRs. In Appendix C, we empirically showed that this statistical filtering approach directly improve the clustering performance on the discovery cohort and the whole data.

The number of variable groups in the variable clustering stage is chosen from {50, 100}. As mentioned earlier, Kolmogrov-Smirnov nonparametric test is applied to compute the absorption ratios. Variable clusters with the absorption ratio above *τ* ∈ {1, 0.05, 0.01, 10^−4^} are eliminated. Note that in case that *τ* = 1 no variable cluster will be dropped. This step is beneficial even though the underlying data has undergone batch correction (as in our case) as it can reduce the amount of residual batch, and assist with the correction even if the batches are not known.

To reduce the dimensionality of data for each variable cluster we used either principal component analysis (PCA) or independent component analysis (ICA). The number of principal components (hidden components) in PCA (ICA) is chosen from the set {1, 5, 20}, which results in a final signal dimension to be effectively less than the initial data dimension. Moreover, the number of groups for the final clustering is set to *K* = 3. Clustering performance for the FCRC using either k-means or GMM is reported in Table 1(a). The results are also compared to a baseline approach which clusters the samples using k-means or GMM directly after filtering the variables using the IQRs. The final representation of clustered data is visualized in Figure 2a (initial data) and 2b (including the validation cohort). The result shows that the three clusters re-capitulate many of the control, Charge, and Kabuki samples, respectively.

**Table 1:** Performance of FCRC on CHARGE-Kabuki, and Weaver-Sotos data

| Data Subset | (a) CHARGE/Kabuki |  |  |  | (b) Weaver/Sotos |  |  |  |
| --- | --- | --- | --- | --- | --- | --- | --- | --- |
|  | Method | NMI | ARI | Purity | Method | NMI | ARI | Purity |
| Discovery Cohort | FCRC(GMM-PCA) | 0.27 ± 0.13 | 0.17 ± 0.07 | 0.60 ± 0.06 | FCRC(GMM-PCA) | 0.66 ± 0.00 | 0.67 ± 0.01 | 0.71 ± 0.01 |
| | FCRC(kmeans-PCA) | 0.26 ± 0.07 | 0.206 ± 0.06 | 0.64 ± 0.04 | FCRC(kmeans-PCA) | 0.66 ± 0.00 | 0.67 ± 0.01 | 0.72 ± 0.01 |
| | FCRC(GMM-ICA) | 0.54 ± 0.02 | 0.46 ± 0.01 | 0.80 ± 0.00 | FCRC(GMM-ICA) | 0.66 ± 0.00 | 0.67 ± 0.01 | 0.71 ± 0.01 |
|  | FCRC(kmeans-ICA) | **0.61 ± 0.02** | **0.48 ± 0.04** | **0.81 ± 0.03** | FCRC(kmeans-ICA) | **0.67 ± 0.01** | **0.68 ± 0.01** | **0.72 ± 0.01** |
| | GMM (baseline) | 0.23 ± 0.14 | 0.19 ± 0.11 | 0.59 ± 0.12 | GMM (baseline) | 0.66 ± 0.00 | 0.67 ± 0.01 | 0.71 ± 0.01 |
| | kmeans (baseline) | 0.23 ± 0.02 | 0.16 ± 0.03 | 0.59 ± 0.03 | kmeans (baseline) | 0.66 ± 0.00 | 0.67 ± 0.01 | 0.71 ± 0.01 |
| Whole Dataset | FCRC(GMM-PCA) | 0.23 ± 0.04 | 0.16 ± 0.05 | 0.50 ± 0.03 | | | | |
| | FCRC(kmeans-PCA) | 0.19 ± 0.02 | 0.14 ± 0.02 | 0.51 ± 0.03 | | | | |
| | FCRC(GMM-ICA) | 0.43 ± 0.09 | 0.32 ± 0.05 | 0.68 ± 0.03 | | | | |
|  | FCRC(kmeans-ICA) | **0.46 ± 0.01** | **0.33 ± 0.01** | **0.70 ± 0.01** |  |  |  |  |
| | GMM (baseline) | 0.15 ± 0.04 | 0.10 ± 0.06 | 0.48 ± 0.04 | | | | |
| | kmeans (baseline) | 0.18 ± 0.01 | 0.13 ± 0.04 | 0.47 ± 0.03 | | | | |

**Figure 2:**
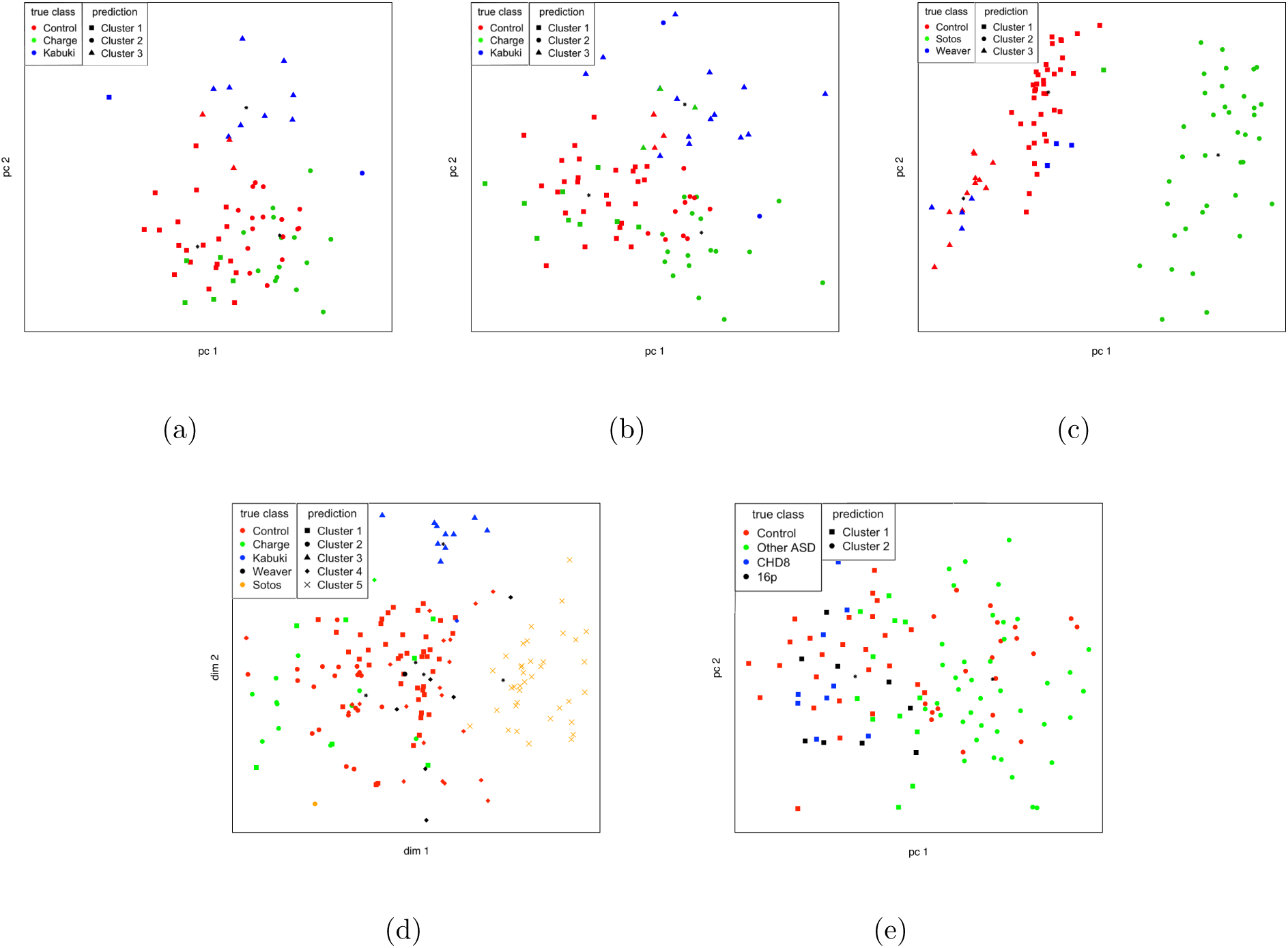
Visualization of the clustering result using the FCRC methods for different datasets: For all the datasets except the combined data the first two principal components of the final signals are used. Due to visualization difficulties of the clustering of combined dataset samples, t-SNE plot is used for the visualization of the combined dataset. (a) The 3 clusters for the discovery cohort of the CHARGE/Kabuki data (b) The 3 clusters for the union of discovery and validation cohort in the CHARGE/Kabuki data. (c) The 3 clusters for the Weaver/Sotos data. (d) The 5 clusters for the combined dataset (CHARGE/Kabuki/Weaver/Sotos). (e) The 2 clusters of the de-novo experiment on distinguishing between unspecified ASD samples from the controls and a set of samples with a specified ASD variant.

### 4.3 Sotos and Weaver Syndromes Data

For the Sotos and Weaver syndromes data [3], we applied the same procedure as with the CHARGE and Kabuki syndromes. To validate the replicability of the method, we did not change any of the hyperparameters of the FCRC procedure. The results is reported in the Table 1(b), and visualized in Figure 2c. *K* = 3 is used as the number of clusters. We note that applying FCRC to Sotos and Weaver data results lower variance clustering than applying it to CHARGE and Kabuki data. This is largely driven by the ease of identification of the Sotos cluster. As previously discussed, the gene responsible for Sotos disorder, NSD1, is a methyltransferase, and significantly changes methylation throughout the genome, creating an unusually strong signature [3].

### 4.4 Combination of the CHARGE/Kabuki and Weaver/Sotos Data

To verify the effectiveness of FCRC on distinguishing more than two diseases, we combined the cohorts of CHARGE/Kabuki and Weaver/Sotos from the previous sections. The sets of used hyperparameters are the same as those of for the CHARGE/Kabuki and Sotos/Weaver dataset clustering, with the exception of requiring 5 groups in the final clustering step. Clustering performance of FCRC is reported in Table 2(a) and in Figure 2d. The performance is comparable to the clustering performance on each of the datasets individually, showing the ability of the model to distinguish amongst a larger number of disorders.

**Table 2:** Performance of FCRC on the the combined data (CHARGE/Kabuki/Weaver/Sotos) and on an orthogonal Autism dataset

| (a) Combined CHARGE/Kabuki/Weaver/Sotos |  |  |  | (b) Autism Variants |  |  |  |
| --- | --- | --- | --- | --- | --- | --- | --- |
| Method | NMI | ARI | Purity | Method | NMI | ARI | Purity |
| FCRC(GMM-PCA) | **0.60 ± 0.04** | 0.45 ± 0.05 | 0.62 ± 0.05 | FCRC(GMM-PCA) | 0.16 ± 0.01 | 0.22 ± 0.02 | 0.74 ± 0.01 |
| FCRC(kmeans-PCA) | 0.45 ± 0.02 | 0.33 ± 0.03 | 0.58 ± 0.04 | FCRC(kmeans-PCA) | **0.19 ± 0.02** | **0.26 ± 0.03** | **0.76 ± 0.02** |
| FCRC(GMM-ICA) | 0.57 ± 0.04 | 0.40 ± 0.05 | 0.55 ± 0.07 | FCRC(GMM-ICA) | 0.11 ± 0.03 | 0.15 ± 0.06 | 0.71 ± 0.04 |
| FCRC(kmeans-ICA) | 0.57 ± 0.05 | **0.47 ± 0.06** | **0.65 ± 0.06** | FCRC(kmeans-ICA) | 0.14 ± 0.02 | 0.18 ± 0.02 | 0.72 ± 0.01 |
| GMM (baseline) | 0.42 ± 0.03 | 0.25 ± 0.06 | 0.47 ± 0.03 | GMM (baseline) | 0.15 ± 0.03 | 0.21 ± 0.03 | 0.74 ± 0.01 |
| kmeans (baseline) | 0.45 ± 0.01 | 0.32 ± 0.05 | 0.54 ± 0.06 | kmeans (baseline) | 0.12 ± 0.01 | 0.18 ± 0.011 | 0.72 ± 0.01 |

### 4.5 Autism Syndromes Data

To further test the model in a setting without a ground truth we applied it to an autism syndromes data (MTS & RW, in preparation) which contained two genetically distinct variants of autism, due to mutations in CHD8, and deletions on chromosome 16p, as well as genetically undifferentiated autism samples and a set of controls. At this proof of concept phase we asked for only two clusters to explain the data. In Table 2(a) we compared the performance of FCRC with the baselines for this task. It is worth noting that slightly worst clustering performance can be obtained using clustering individuals directly applying the final clustering on the data obtained after the second step of the FCRC (Figure 1). Even though the full FCRC procedure has more steps, it can increase the clustering performance by removing suspicious signals which increases the generalization of FCRC to be used for a similar data as well.

The final clustering result revealed a cluster highly enriched in autism samples (on the right of Figure 2e). This cluster contained 44/52 undifferntiated ASD samples and only 15/48 controls, a statistically significant enrichment (p-value of < .00001 chi-squared test). We also checked whether this enrichment was due to batch; the cluster contained 64/102 samples for one specific batch. Nevertheless, the enrichment of ASD samples relative to the batch is still statistically significant (p-value of .005042).

## 5 Discussion

To enable the unsupervised identification of diseases without a known DNAm signature, we propose FCRC, an unsupervised learning method which we applied on several DNA methylation array datasets. Our results show that the method can differentiate between known human disorders without *a priori* signatures, and shows promise for analysis of a more common disease (ASD).

In our experiments, we observed that further tuning the parameters in the FCRC procedure could improve the performance. Recent development of hyperparameter optimization and metalearning approaches could be helpful in determining better performing hyperparameters. Moreover, as mentioned earlier, using other preprocessing techniques can increase the statistical power of the proposed method and reduce the performance variation. Therefore, a straightforward continuation of this work might be to use these preprocessing methods to get a better performance.

The success of deep learning in the areas such as natural language processing, computer vision, generative modeling, and reinforcement learning motivates researchers to leverage deep neural networks in computational biology. Several works have successfully applied deep models for regulatory genomics and biological image data [23]. DNA methylation array data typically consist of a few samples. Using deep neural networks for this type of data is not only challenging because of the sample size, but also due to the particular noises and widespread biases such as batch effect, which is an active area of research. Moreover, building a deep model that directly learns from biological samples as vectors with several hundred thousand dimensions is another challenge. Using an appropriate feature selection method, increasing the data quality, and combining multiple datasets are options that could help alleviate the existing challenges.

## Acknowledgements

The authors would like to thank David Duvenaud, Shadi Zabad, Marta Skreta, Parisa Shooshtari, and Tahmid Mehdi for helpful discussions and providing useful comments.

## 6 Appendix

### A Detailed FCRC

The practical FCRC procedure that we have used in our experiments is shown in algorithm 2

#### Algorithm 2 Implemented FCRC procedure for DNA methylation data

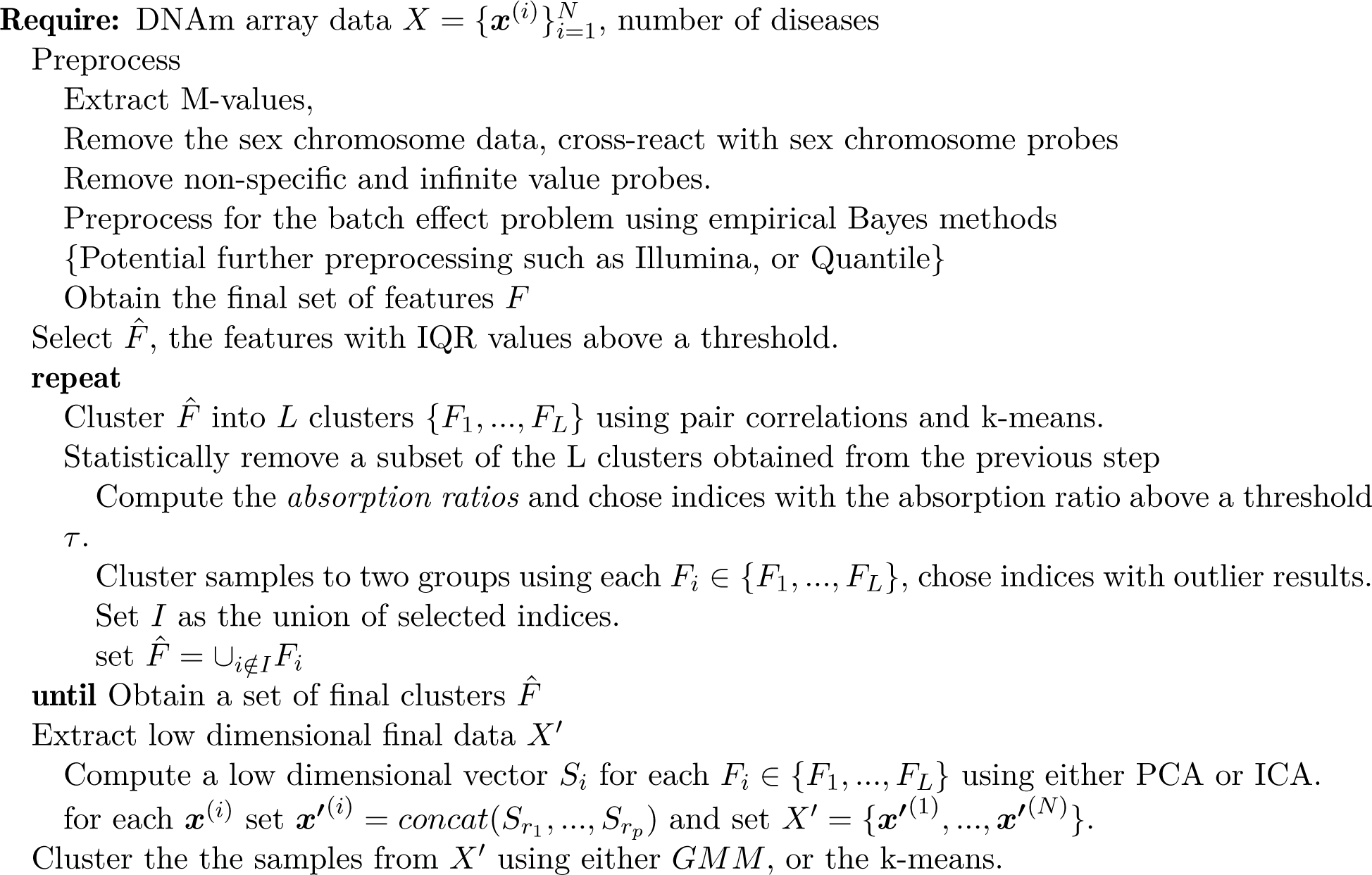

### B Potential negative effect of nearest neighbor graph based feature selection methods for DNAm array data

There are various supervised and unsupervised feature selection methods that are based on local structure of the data using nearest neighbors of data points. For example, Laplacian score [24], MCFS [16], or SPEC [25] use the nearest neighbor graph built from inputs distance matrix. In this section, using Euclidean distance, we show the potential negative effects of leveraging nearest neighbor feature selection methods in further downstream analysis in cases that we have batch effect problem.

As a real example, we used the Charge and Kabuki data [2]. For *k* ∈ {1, …, *N* − 1}, where N is the total number of individuals, we compute the empirical probability that one of *k* nearest neighbors of an individual falls in the same batch. We expect that this probability should vary much over different values of *k*. Figure 3(a) shows the value of this empirical probability, before and after the batch effect processing used in section 4.2. As we can see, the batch effect processing effectively decreases the mentioned empirical probability, at least for lower *k* values.

**Figure 3:**
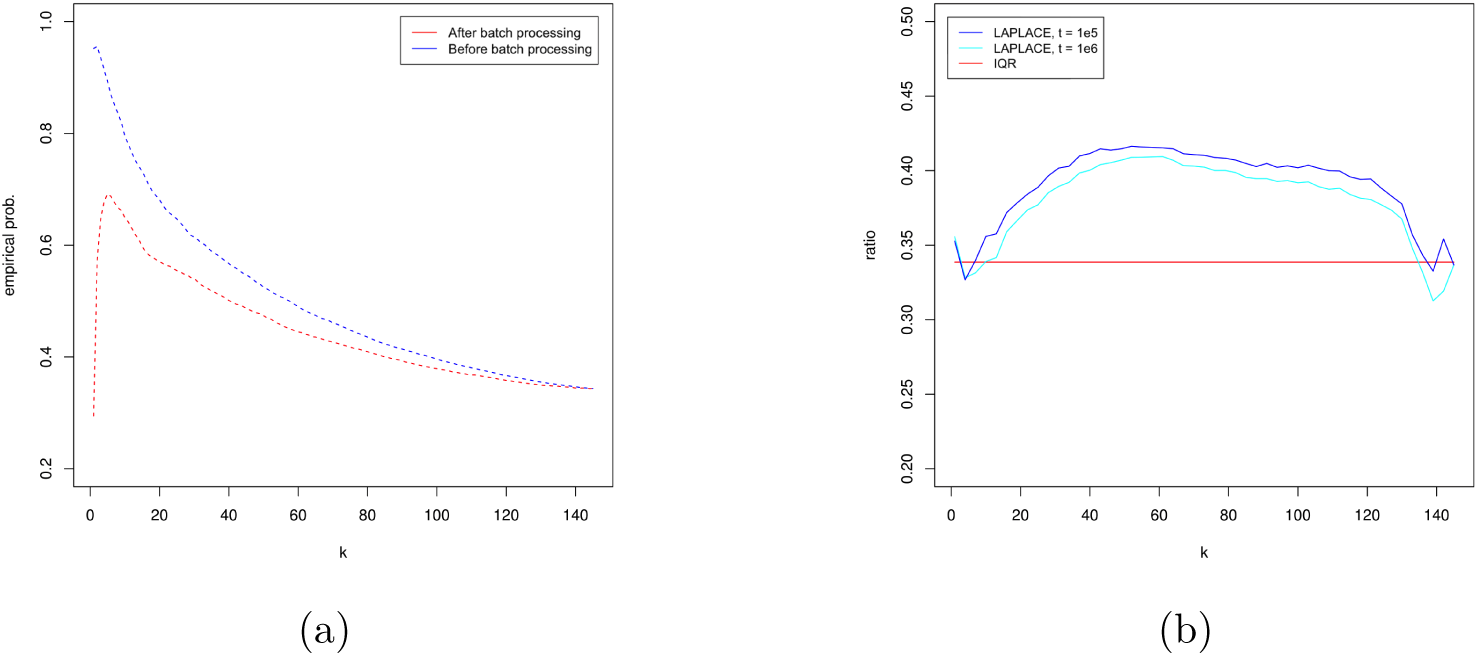
(a): Empirical probability that an individual selects another individual within the same batch as one of its *k* nearest neighbors before and after batch problem processing. (b) ratio of the feature that still suffer from batch effect after batch processing to the total number of filtered features chosen by simple IQR feature filtering, and Laplacian score feature selection with the exponent parameter *t*.

To view the potential negative effect, we compare the Laplacian score feature selection, and the simple IQR filtering we used in our experiments. After processing batches using the mentioned method, we ran t-test for the values of each feature over pair of batches. We labeled the features that their test results had at least one p-value less than 0.005 as features that still suffer from batch effect problem. In figure 3(b) we see the ratio of the labeled features to the total number of labeled features in the set of 20*k* filtered features, obtained from Laplacian score and IQR feature selection approaches over different values of *k*. As we can see, for most values of *k* the simple IQR feature filtering which is independent of *k* performs better in terms of removing the batch affected features than the Laplacian feature selection procedure. Therefore, we argue that *k* must be chosen very carefully in order to avoid further bias in downstream analysis.

### C IQR as a useful statistic for filtering features

In this section we provide a simple and empirical study that the IQR is a useful statistic to filter features. For our experiment, we chose Charge-Kabuki dataset and sorted the features based on their IQR values. We clustered data based on the first top *s* features, second top *s* features, and so on. We measured the NMI value to compare the clustering result with the ground truth. The results is shown in figure 4. It is clear in the graphs that the NMI values is positively correlated with the group IQR values, which means filtering based on IQR could potentially increases the ratio of the useful features for clustering.

**Figure 4:**
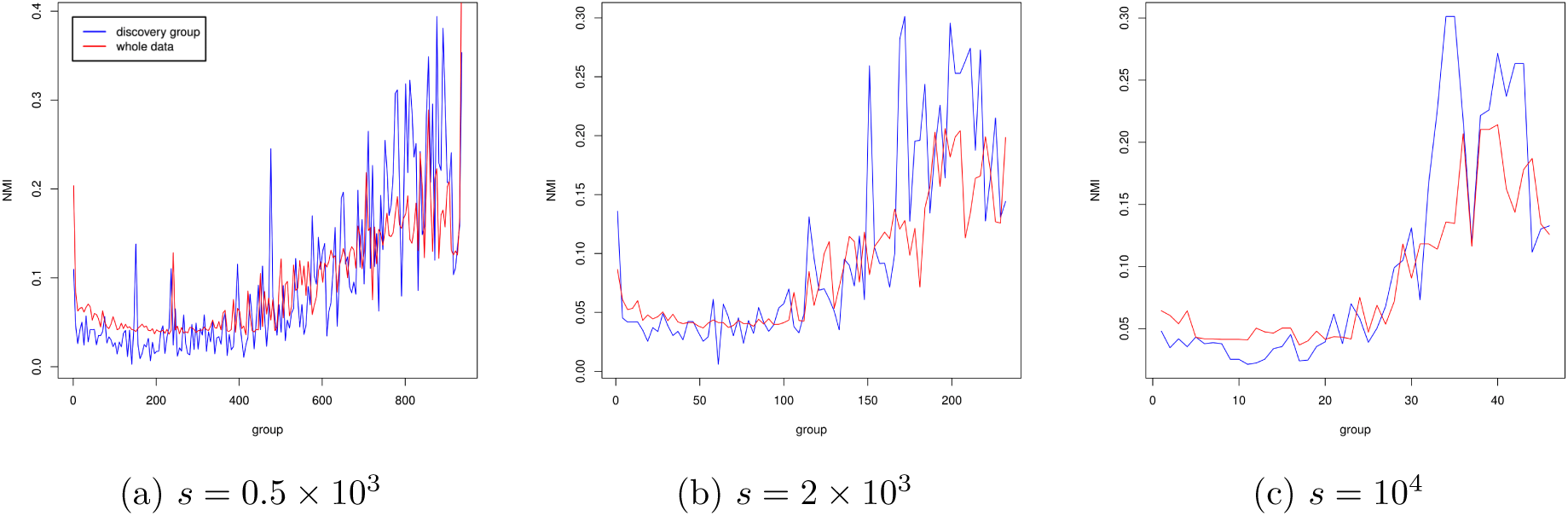
The NMI score as a result of clustering the individuals in the Charge-Kabuki data based on a group of features. As the group number on the horizontal axis increases the value of feature IQRs within the group increases. *s* specifies the number of features within the group.

### D Sample confusion matrices

Confusion matrices of a sample clustering for each data using the FCRC method are reported in the table 3. Values in the parentheses represent the equivalent clustering confusion matrices using the baseline approach.

**Table 3:** Confusion Matrices

| (a) Charge-Kabuki (Discovery Cohort) |  |  |  |  | (b) Charge-Kabuki (Whole Data) |  |  |  |  |  |  |  |
| --- | --- | --- | --- | --- | --- | --- | --- | --- | --- | --- | --- | --- |
|  |  | Reference |  |  |  |  | Reference |  |  |  |  |  |
|  |  | Control | Charge | Kabuki |  |  | Control | Charge | Kabuki |  |  |  |
| Prediction | Cluster 1 | 29 (23) | 2 (2) | 0 (6) | Prediction | Cluster 1 | 28 (29) | 13 (5) | 1 (8) |  |  |  |
|  | Cluster 2 | 10 (8) | 16 (15) | 1 (4) |  | Cluster 2 | 10 (11) | 24 (18) | 2 (7) |  |  |  |
|  | Cluster 3 | 1 (9) | 1 (2) | 10 (1) |  | Cluster 3 | 2 (0) | 2 (16) | 16 (4) |  |  |  |
| (c) Sotos-Weaver |  |  |  |  | (d) Autism Syndromes |  |  |  |  |  |  |  |
|  |  | Reference |  |  |  |  | Reference |  |  |  |  |  |
|  |  | Control | Sotos | Weaver |  |  | Control | Other ASDs | CHD8 | 16p | 16p sig. | CHD8 sig. |
| Prediction | Cluster 1 | 38 (38) | 1 (1) | 3 (3) | Prediction | Cluster 1 | 33 (34) | 15 (20) | 9 (9) | 9 (9) | 9 (9) | 7 (7) |
|  | Cluster 2 | 0 (0) | 37 (37) | 0 (0) |  | Cluster 2 | 15 (14) | 37 (32) | 0 (0) | 0 (0) | 0 (0) | 0 (0) |
|  | Cluster 3 | 12 (12) | 0 (0) | 4 (4) |  |  |  |  |  |  |  |  |
| (e) Combined dataset |  |  |  |  |  |  |  |  |  |  |  |  |
|  |  | Reference |  |  |  |  |  |  |  |  |  |  |
|  |  | Control | Charge | Kabuki | Weaver | Sotos |  |  |  |  |  |  |
| Prediction | Cluster 1 | 47 (36) | 5 (0) | 1 (3) | 2 (2) | 0 (1) |  |  |  |  |  |  |
|  | Cluster 2 | 19 (20) | 9 (14) | 1 (3) | 0 (0) | 0 (0) |  |  |  |  |  |  |
|  | Cluster 3 | 0 (0) | 4 (3) | 8 (1) | 0 (1) | 4 (0) |  |  |  |  |  |  |
|  | Cluster 4 | 24 (19) | 1 (2) | 0 (4) | 5 (4) | 0 (0) |  |  |  |  |  |  |
|  | Cluster 5 | 0 (15) | 0 (0) | 0 (0) | 0 (0) | 34 (37) |  |  |  |  |  |  |

### E PCA plot for the combined dataset

For further clarification, the PCA plot for the clustering result of the combined dataset with and without the controls is provided in the figure 5.

**Figure 5:**
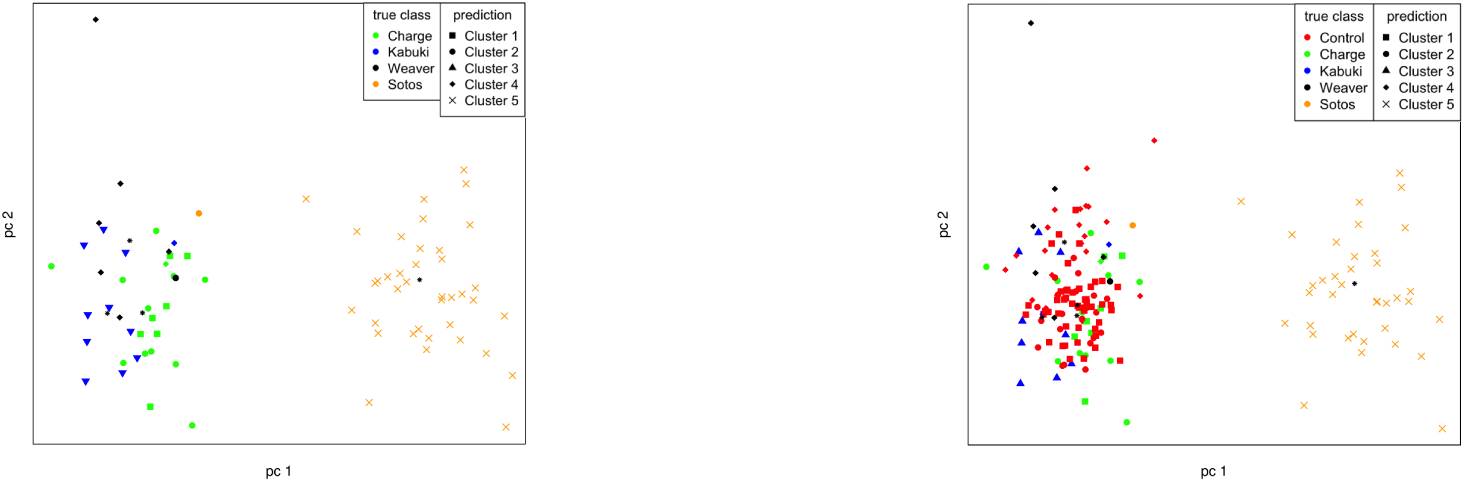
The PCA plot for the clustering result of the combined data

